# A Deep Denoising Sound Coding Strategy for Cochlear Implants

**DOI:** 10.1101/2022.11.11.516123

**Authors:** Tom Gajecki, Yichi Zhang, Waldo Nogueira

## Abstract

Cochlear implants (CIs) have proven to be successful at restoring the sensation of hearing in people who suffer from profound sensorineural hearing loss. CI users generally achieve good speech understanding in quiet acoustic conditions. However, their ability to understand speech degrades drastically when background interfering noise is present. To address this problem, current CI systems are delivered with front-end speech enhancement modules that can aid the listener in noisy environments. However, these only perform well under certain noisy conditions, leaving quite some room for improvement in more challenging circumstances. In this work, we propose replacing the CI sound coding strategy with a deep neural network (DNN) that performs end-to-end speech denoising by taking the raw audio as input and providing a denoised electrodogram, i.e., the electrical stimulation patterns applied to the electrodes across time. We specifically introduce a DNN that emulates a common CI sound coding strategy, the advanced combination encoder (ACE). We refer to the proposed algorithm as ‘Deep ACE’. Deep ACE is designed not only to accurately code the acoustic signals in the same way that ACE would but also to automatically remove unwanted interfering noises, without sacrificing processing latency. The model was optimized using a CI-specific loss function and evaluated using objective measures as well as listening tests in CI participants. Results show that, based on objective measures, the proposed model achieved higher scores when compared to the baseline algorithms. Also, the proposed deep learning-based sound coding strategy gave eight CI users the highest speech intelligibility results.

## I. Introduction

A cochlear implant (CI) is a surgically implanted neuroprosthetic device that restores the sensation of hearing in people who suffer from profound sensorineural hearing loss. The CI sound coding strategy is responsible for computing the electric stimulation current levels from the audio captured by the CI sound processors’ microphone. There are several CI sound coding strategies used in the industry [1]. Out of these, a widely used sound coding strategy is the continuous interleaved sampling (CIS) [2]. CIS decomposes the incoming sound into multiple different frequency bands, which are used to modulate electric pulses that stimulate the auditory nerve. The set of pulses is sent to the available electrodes to stimulate the auditory nerve across time in an interleaved way. Other strategies perform band selection by picking the most perceptually relevant bands for stimulation.

Band selection has the advantage of reducing power consumption without compromising speech intelligibility, which is the reason why it is widely used in the CI industry. Some common criteria to select relevant bands are based on magnitude, used in the advanced combination encoder (ACE) [3], or on psychoacoustic masking, used in the PACE/MP3000 sound coding strategy [4]. When these CI sound coding strategies are used, the electrodes located near the base of the cochlea represent higher frequencies, whereas those located in the most apical region transmit low-frequency information. In this work, we focus specifically on the ACE sound coding strategy. However, the presented approach could be generalized to any available sound coding strategy, as all of them generate electrodograms (i.e., the normalized amplitudes that are subsequently mapped to the current levels that each electrode will deliver to the auditory nerve over time).

In general, a CI together with its corresponding sound coding strategy allows the user to understand speech in quiet conditions, however, it fails to do so when loud interfering signals (i.e., at low signal-to-noise ratios; SNRs), such as noise or other talkers, are present [5]. In order to overcome the limitations that CI users face in noisy conditions, many speech enhancement techniques have been proposed to improve speech intelligibility, such as spectral contrast enhancement [6], [7], spectral subtraction [8], Wiener filtering [9] and time-frequency masking [10]. Although these techniques work reasonably well, recently the signal processing community has been leaning towards more modern data-driven approaches to perform single-channel speech enhancement, such as deep learning models [11]–[14].

Modern approaches to source separation and speech enhancement typically utilize time-frequency representations of the input signals for extracting features, which can lead to highly effective results [15], [16]. However, these do not exploit potentially rich sources of information, such as the phase, limiting speech separation quality. To overcome this problem, end-to-end deep learning-based approaches that directly work in the time domain have been recently proposed. For example, [17] proposed a fully-convolutional time-domain audio separation network (Conv-TasNet), a deep learning framework for end-to-end time-domain speech separation. This model addresses the shortcomings of separation in the frequency domain, achieves state-of-the-art performance, and is suitable for low-latency applications. Thus, approaches that perform end-to-end processing are getting more attention in the community, making them an attractive potential solution to the CI ‘cocktail party’ problem [18]. A front-end approach, however, may not fully exploit the CI processing characteristics.

In order to optimize speech enhancement for CIs, it may be beneficial to design algorithms that consider the CI processing scheme. Hence, there has been some work done specifically for CIs, where DNNs are included in the CI signal path [14], [19]. These approaches reduce noise, for example, by directly applying masks in the filter bank used by the CI sound coding strategy. Recently, inspired by the aforementioned Conv-TasNet, [20] proposed a deep learning-based end-to-end CI sound coding strategy, referred to as ‘Deep ACE’.

Deep ACE replaces the clinical ACE sound coding method and automatically performs speech enhancement by estimating denoised electrodograms directly from raw audio. It leverages audio-to-electrodogram domain transformation to improve noise reduction for CIs. Although phase information is not necessary for the synthesis of the electrodograms, using it may help generate a proper input signal encoding, and ergo, a better latent representation. Deep ACE is intended to take advantage of such signal representation in order to extract global patterns from its characteristics, identifying which ones are more likely to be embedding speech content.

This study extensively examines Deep ACE [20], introducing a novel and improved topology, along with an optimized hyperparameter configuration that enhances the model’s generalization capabilities. The model was trained on a large dataset and optimized through a loss function tailored for CI listening that discourages the activation of irrelevant bands, with the aim of improving speech comprehension for CI users. The study evaluates the proposed model and compares it with baseline algorithms using objective measures and listening tests with CI users to determine if Deep ACE outperforms the tested baselines and the existing clinical ACE sound coding strategy.

## II. Methods & Materials

### A. Advanced combination encoder (ACE)

The ACE sound coding strategy processes the acoustic signal captured by the microphone, by first sampling it at 16 kHz. Then, a filter bank implemented as a 128-point fast Fourier transform (FFT), commonly with a 32-point hop size, is applied, introducing a 2 ms algorithmic latency (this will depend on the channel stimulation rate; CSR). Next, an estimation of the desired envelope is calculated for each spectral band *E_k_*, (*k* = 1,…, *M*). Each spectral band is mapped to an electrode and represents one channel. *M* denotes the total number of channels/electrodes. In this study, the band selection block sets *N* = 8 out of *M* = 22 envelopes by selecting the ones with the largest amplitudes, which are then non-linearly compressed by a loudness growth function (LGF) given by:

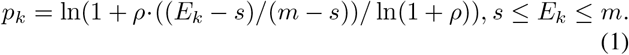

The output of the LGF at band *k* (*p_k_*) represents the normalized stimulation amplitude used to stimulate the auditory nerve using electrode *k*. The stimulation patterns across electrodes obtained from the LGF output over time constitute the electrodogram. For values of *E_k_* below base level *s, p_k_* is set to zero, and for values of *E_k_* above saturation level *m, p_k_* is set to one. We used *ρ* = 416.2, *s* = 4/256, and *m* = 150/256 in our experiments.

Finally, the last stage of the sound coding strategy maps every *p_k_* into the subject’s dynamic range between threshold levels and most comfortable levels for electrical stimulation. The *N* selected electrodes are stimulated sequentially for each audio frame, representing one stimulation cycle. The number of cycles per second thus determines the CSR. A block diagram showing the described processes is shown in Figure 1a; ACE.

**Fig. 1.**
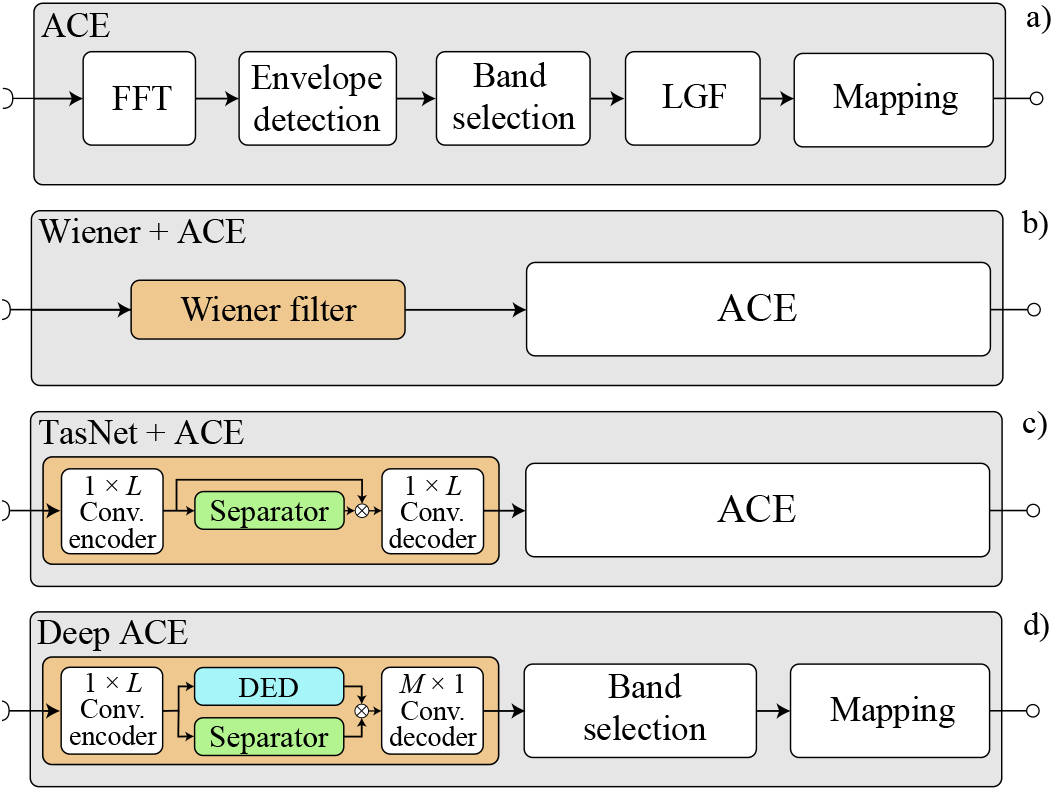
Block diagrams of the four different signal processing systems. In c) and d) *L* refers to the length of the filters used to encode the input signal (and to decode it in the case of TasNet+ACE) and *M* to the number of available CI spectral bands. White circles at the output represent the CI external coil.

### B. Speech enhancement algorithms

#### 1) Wiener filter (baseline #1)

Here, we use a classic front-end signal processing method based on Wiener filtering, a widely used technique for speech denoising that relies on a priori SNR estimation [21] (Figure 1b; Wiener+ACE). Different variations of this algorithm are used in commercially available single-channel noise reduction systems included in CIs [22], [23]. Therefore, this classic algorithm is an appropriate baseline to use when developing new speech enhancement methods in the context of CIs [19].

#### 2) Conv-TasNet (baseline #2)

The front-end DNN-based baseline system used in this study is the well-known conv-TasNet (which we will refer to as TasNet for simplicity) [17]. This system performs end-to-end audio speech enhancement and feeds the denoised signal to ACE, where further processing is performed to obtain the electrodograms (Figure 1c; TasNet + ACE). The TasNet structure has proven to be highly successful for single-speaker speech enhancement tasks, improving state-of-the-art algorithms, and obtaining the highest gains with modulated noise sources [24].

#### 3) Deep ACE (proposed method)

This architecture builds upon the previously developed deep denoising sound coding strategy described in [20]. Deep ACE is designed to estimate the output of the LGF by taking in raw audio input and predicting the denoised electrodograms. This approach is independent of individual CI fitting parameters and maintains the standard ACE strategy’s 2 ms total algorithmic delay. The enhancer module in the here presented Deep ACE contains three main differences when compared to the one in TasNet+ACE (Figure 1d; Deep ACE). The previous version of the model presented in [20] shared two of these dissimilarities with the current version. These were the use of a trainable antirectifier unit as the activation function in the encoder and the output dimensionalities at the decoder. For details, refer to [20].

The primary architectural innovations in the here presented Deep ACE, are the inclusion of a deep envelope detector (DED) positioned in the skipping path of the original Deep ACE model [20], and the improvement of hyperparameter configuration. The DED replaces the envelope detection block in the original ACE (see the ‘DED’ block in Figure 1d). This module performs dimensionality reduction between the encoder and the decoder modules to match the number of bands to be stimulated and to extract other essential features from the encoded signal. This process is necessary for implementation purposes, specifically for the employed loss function (refer to Section II-B5, eq. 5), and it involves three consecutive 1-D convolution layers that are stacked together. The code for training and evaluating Deep ACE can be found online^1^.

#### 4) Model training setup

The deep learning models were trained using batches of two audio segments, each lasting for a duration of 4 seconds, and were trained for a maximum of 100 epochs. In order to achieve optimal results, the initial learning rate was set to 1e-3, which was subsequently reduced by half if the validation set’s accuracy did not show any improvement during three consecutive epochs. To further regularize training, early stopping with 5-epoch patience was applied. Finally, only the best-performing model was saved after the training session.

To optimize the different models, we used the Adam first-order gradient-based optimization algorithm for stochastic objective functions [25]. The utilized range of hyperparameters is presented in detail in Table I. Note that the hyperparameters used in this work have been adjusted through empirical testing to improve the overall models’ performance when compared to the ones used in [20]. For a comprehensive description of the various parameters, interested readers can refer to [17].

**TABLE I.** Hyperparameters used to train the deep learning models

| Parameter | TasNet | Deep ACE |
| --- | --- | --- |
| N | 64 | 64 |
| L | 32 | 32 |
| B | 64 | 64 |
| H | 128 | 128 |
| S | 32 | 32 |
| P | 3 | 3 |
| X | 8 | 8 |
| R | 3 | 3 |
| Model size | 526k | 552k |
| Loss function | SI-SNR (eq. 2) | $\mathcal{L}_\delta$ (eq. 5) |
| Output domain | Audio | Electrodeogram |

#### 5) Model training objectives

In the case of the TasNet+ACE algorithm, the optimizer was used to maximize the scale-invariant (SI) SNR [26] at the output of the TasNet, before being processed by the ACE sound coding strategy (see Figure 1c). The SI-SNR between a given signal with *T* samples, 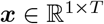 and its estimate 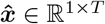 is defined in eq. 2.

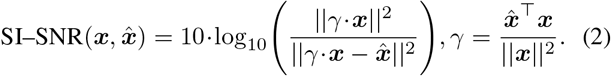

In the Deep ACE model, the decoder module is developed to predict the output at the LGF of ACE to be fed into the band selection process. Therefore, the cost function employed to train it will be based on the mean-squared error, denoted by 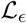. The 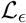 between an *M*-channel and *F*-frame target LGF output, 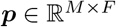 and its estimate 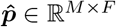, is defined as:

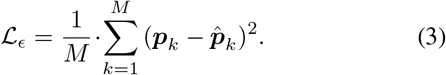

In this work, Deep ACE is optimized by minimizing a variant of the loss function used in [20] (i.e., 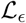). Specifically, we combine the loss function defined in eq. 3 with a punishment term that aims at removing CI stimulation in unwanted channels. Specifically, to penalize we introduce a second loss term that is measured by means of the binary cross entropy between the ideal target mask 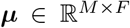 and the estimated mask 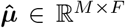 (at the output of the separator). We will denote this function by 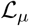, and its value computed in NATS is given by:

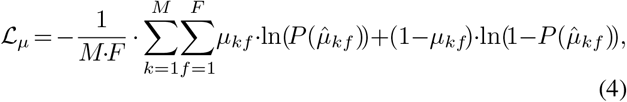

where *μ_kf_* is equal to one if channel *k* at frame *f* contains speech, and to zero otherwise (also known as an ideal binary mask). 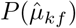 is the predicted probability that channel *k* in frame *f* contains speech. The cost function used to optimize Deep ACE is denoted as 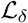, and was constructed by linearly combining 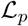 and 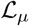 as follows:

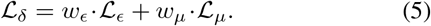

Empirical testing was used to determine the values for the multiplicative weighting factors *w_ϵ_* and *w_μ_*, which were then established as 15 and 1, respectively. The basis for this cost function is rooted in prior research [27], which demonstrated that individuals using CIs can withstand significant distortions in speech segments provided that the selection of frequency bands is accurate.

It is important to note that the second loss term is applied at the separator output, which means that the estimated mask must have the same dimensions as the LGF output. To achieve this, Deep ACE utilizes a DED module (described in II-B3) in the skipping path to decrease the channel dimension of the encoded input and enable the masking operation (see Figure 1d). In addition, the motivation for developing this module is linked to the fact that it is also a component within the ACE sound coding strategy. In a similar manner, it is responsible for minimizing the dimensionality between the filter bank (FFT) and the band selection block (as depicted in Figure 1a). Specifically, the envelope detector in ACE consolidates the frequency bins obtained from the spectral transformation into the number of available electrodes (*M*).

**Fig. 2.**
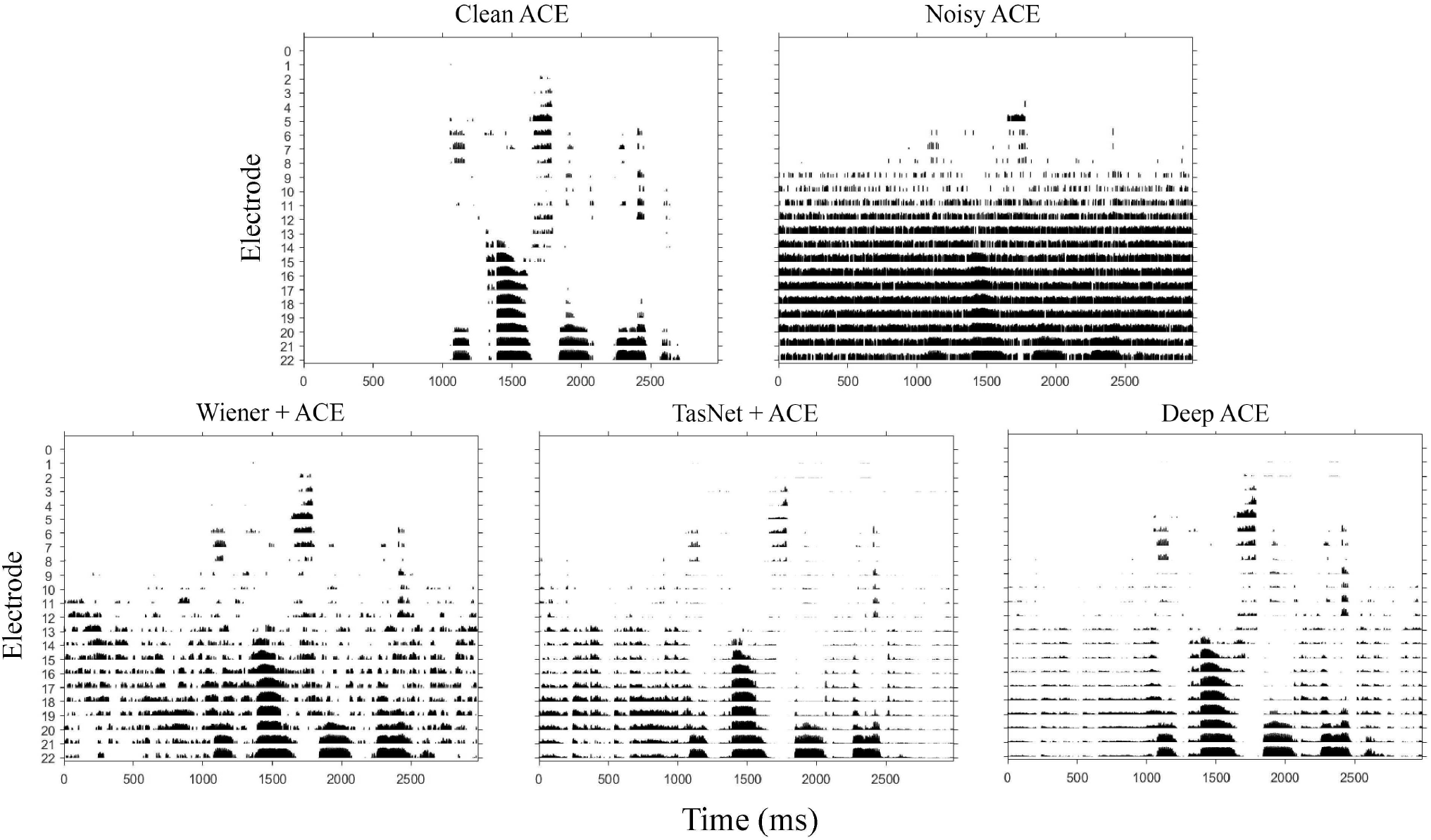
Electodograms after the band selection and mapping processing blocks for the clean, noisy, and processed noisy speech signals processed by the different algorithms. Electrode numbers increase as the mapped frequencies decrease.

### C. Audio material

In this work, we used a total of three different speech datasets and three noise types to assess the models’ performance and generalization abilities. All these audio sets will be described in this section. As a preprocessing stage, all audio material was set to mono and re-sampled at 16 kHz. The corresponding electrodograms were obtained by processing all audio data with the ACE sound coding strategy at an output channel stimulation rate of 1,000 pulses per second CSR.

#### 1) Speech data

##### a) LibriVox corpus [28]

This speech data was originally designed for end-to-end speech translation, however, in this study, we mix the speech material with noise to train our models for speech denoising. The speech data contained in this corpus consists of fluent spoken sentences with a total duration of 18 hours. The quality of audio and sentence alignments was checked by a manual evaluation, showing that speech alignment is in general very high. In fact, the sentence alignment quality is comparable to well-used parallel translation data.

##### b) TIMIT corpus [29]

This corpus contains broadband recordings of 630 people speaking the eight major dialects of American English, each reading ten phonetically-rich sentences. In this work, files from 112 male and 56 female speakers in the test set were selected.

##### c) HSM corpus [30]

Speech intelligibility in quiet and in noise was measured by means of the Hochmair, Schulz, Moser (HSM) sentence test, based on a dataset composed of 30 lists with 20 everyday sentences each (106 words per list).

#### 2) Noise data

##### a) Environmental noises; DEMAND [31]

The environmental noises recorded to create this dataset are split into six categories; four are indoor noises and the other two are outdoor recordings. The indoor environments are further divided into domestic, office, public, and transportation; the open-air environments are divided into streets and nature. There are 3 environment recordings per category.

##### b) Synthetic noises; SSN [32] and ICRA7 [33]

To evaluate the different algorithms, in this work we also use stationary speech-shaped noise (SSN) and non-stationary modulated seven-speaker babble noise (ICRA7) as synthetic interferers.

#### 3) Training, evaluation and testing data

The training set was composed of speech from the LibriVox corpus and noise from the DEMAND dataset. Specifically, 30 male (M) and female (F) speakers were randomly selected from the speech corpus, and two environments were randomly selected from each of the noise categories. For validation, 20% of the training data was used. The noise and speech subsets used for training will be referred to as EN_1_ and LibriVox_1_, respectively. For testing, the remaining audio data was used, namely the noise and two speech sets which will be referred to as EN_2_, LibriVox2, and HSM, respectively.

Speech and noise signals were mixed at SNR values ranging uniformly from −5 to 10 dB. The processed clean speech signals were also included in the listening experiments to assess whether the proposed model introduced perceptually relevant distortions. A description of how the different datasets were distributed for the experiments is shown in Table II.

**TABLE II.**
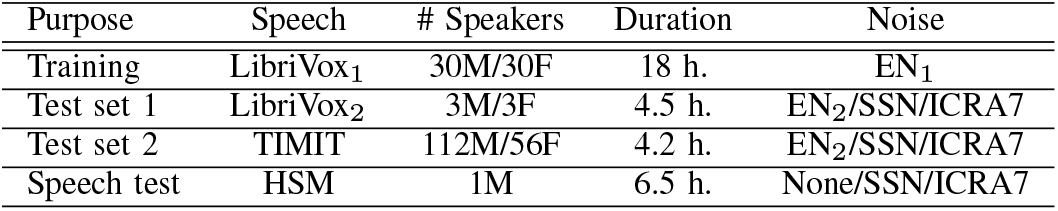
Datasets used to train, validate and test the models

### D. Evaluation

#### 1) Objective evaluation

To assess the objective performance of each of the tested algorithms we compute the amount of noise reduction achieved, electrode-wise correlation coefficients between the denoised and clean signals, and a speech intelligibility score based on the short-time objective intelligibility (STOI) index [34]. Note that in this work we investigate end-to-end CI processing, so the latter objective measure is computed from the synthesized electrodograms (*p*) obtained using a vocoder, resulting in the STOI version used in this work, the vocoder STOI (VSTOI; [35], [36]).

##### a) SNRi

To assess the amount of noise reduction performed by each of the tested algorithms we compute the SNR improvement (SNRi). This measure is calculated in the electrodogram domain and compares the original input SNR to the one obtained after denoising, and is given by:

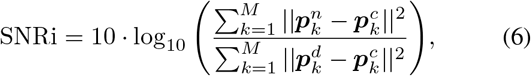

where *p_k_* represents the LGF output of band *k* and the superscripts *n, c*, and *d* are used to denote the noisy, clean, and denoised electrodograms, respectively.

##### b) LCC

To characterize potential distortions and artifacts introduced by the tested algorithms, the linear correlation coefficients (LCCs) between the clean ACE electrodograms (*p^c^*) and the denoised electrodograms (*p^d^*) were computed. The LCCs were computed channel-wise (i.e., one correlation coefficient was computed for each of the 22 channels) to assess channel output degradation caused by the denoising process. The LCC_*k*_ for band *k* is computed based on the Pearson correlation coefficient [37] as follows:

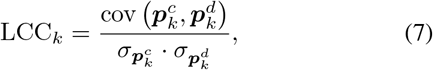

where cov(*X, Y*) is the covariance between *X* and *Y*, and *σ_p_k__* is the standard deviation of the values in the corresponding electrodogram *p_k_*.

##### c) VSTOI

To estimate the speech intelligibility performance expected from each of the algorithms, the VSTOI score [35], [36], [38] was used. This metric relies directly on STOI [34], which is modeled based on normal hearing speech performance. However, VSTOI has proven to be useful in CI studies in order to compare relative expected speech intelligibility outcomes [36]. Specifically, the purpose of this metric is to evaluate the potential relative variations in speech performance that could be achieved in behavioral experiments, rather than providing an exact estimation of an individual’s CI performance. The VSTOI score ranges from 0 to 1, where the higher score represents a predicted higher speech performance.

In this work, speech recognition performance was estimated using the clean unprocessed speech as a reference and the vocoded denoised speech as the processed signal. The vocoded speech was obtained from the electrodograms (*p_k_*) by expanding the amplitudes contained in the electrodogram signals through the inverse LGF operation. Next, the expanded amplitudes contained in each band were used to amplitude-modulate band-pass filtered noise channels. The center frequencies of the band-pass filters used to obtain the modulated noise bands correspond to the ones mapped to each of the CI electrodes. Finally, by summing up all amplitude-modulated noise bands the vocoded signal is obtained.

#### 2) Behavioral evaluation

##### a) Participant demographics

Eight postlingually deafened CI users participated in the listening tests. All participants were native German speakers and had been implanted for several years. They were invited to participate in a 3-hour test at the German Hearing Center of the Hannover Medical School (MHH), for which the travel costs were covered. The experiment was granted ethical approval by the MHH ethics commission. A synopsis of the patient-related data is shown in Table III.

**TABLE III.** CI participant information and experiment settings

| ID | Age | Gender | Tested side | SSN SNR | ICRA7 SNR |
| --- | --- | --- | --- | --- | --- |
| BI01 | 70 | M | R | 0 | 5 |
| BI02 | 40 | M | R | 0 | 5 |
| BI03 | 74 | M | R | 0 | 0 |
| BI04 | 70 | M | R | 5 | 5 |
| BI05 | 36 | F | R | 0 | 0 |
| BI06 | 62 | M | R | 5 | 5 |
| BI07 | 86 | M | R | 10 | 10 |
| BI08 | 57 | F | R | 5 | 5 |

##### b) Experimental setup

The testing material was processed to obtain electrodograms, which were then delivered to the cochlear implant (CI) located in the participants’ selfreported best-performing hearing side (as indicated in Table III) through direct stimulation using the RF GeneratorXS interface (Cochlear Ltd.), controlled by MATLAB and the Nucleus Implant Communicator V.3 (Cochlear Ltd.). The experiments were conducted on a personal computer with custom-made software written in MATLAB. Prior to the commencement of experiments, a hardware security check was conducted by analyzing the generated signals by the research interface with an oscilloscope.

During the experiment, the CI participant was accompanied by two observers in the laboratory. One observer operated the software, while the other counted the number of correctly identified words by marking them on a corresponding printed list. Each listening condition was evaluated twice, using different randomly selected lists of HSM sentences. The final score was computed by averaging the number of correctly identified words for each condition, resulting in the word recognition score (WRS). The test SNR was adjusted to a level where the participant could understand between 20% and 80% of the presented words using the unprocessed ACE noisy condition, and was assessed for the two different types of noise (as shown in Table III).

## III. Results

### 1) Objective instrumental results

#### a) SNRi

Figure 3 illustrates the SNRi obtained with each of the algorithms. The Deep ACE model demonstrated superior performance over TasNet+ACE and Wiener+ACE in all conditions, particularly at low SNRs. This finding suggests that the Deep ACE sound coding approach presented here represents an improvement over the model introduced in [20], where no improvement in SNR was observed compared to the competing front-end deep-learning baseline (TasNet). Moreover, although the SNRi values for the TIMIT and LibriVox_2_ speech datasets were analyzed, they are not reported in this study, but similar patterns were identified under all testing noise conditions. In general, these observations demonstrate a substantial improvement with respect to the previous version of the model presented in [20].

**Fig. 3.**
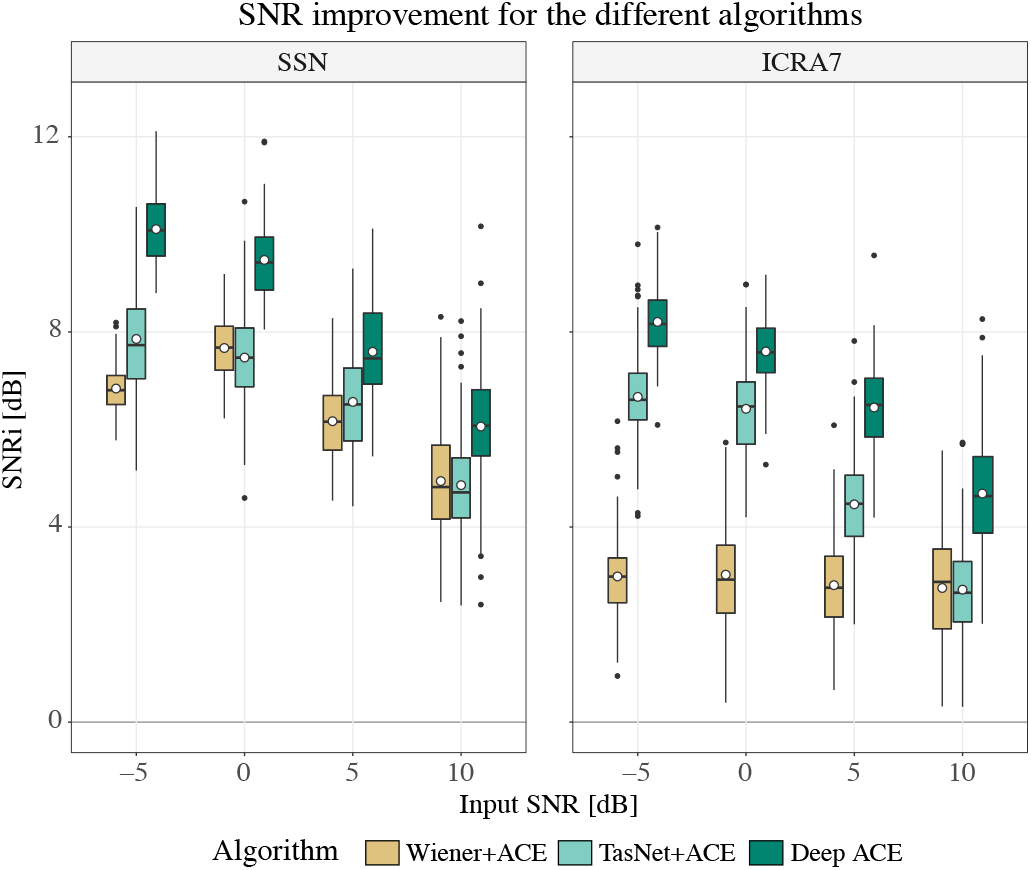
Box plots showing the SNRi scores in dB for the tested algorithms in SSN and ICRA7 noises for the different SNRs using the HSM speech dataset. All pair-wise differences were statistically significant. The black horizontal bars within each box represent the median for each condition, the circle-shaped marks indicate the mean improvement, and the top and bottom extremes of the boxes indicate the *Q*_3_ = 75% and *Q*_1_ = 25% quartiles, respectively. The box length is given by the interquartile range (IQR), used to define the whiskers that show the variability of the data above the upper and lower quartiles (the upper whisker is given by *Q*_3_ + 1.5·IQR and the lower whisker is given by *Q*_1_ – 1.5·IQR [39]). Black dots indicate observations that fall beyond the whisker range (outliers).

#### b) LCC

Here we assess the similarity between the original clean and denoised electrodograms produced by the different algorithms. Figure 4 shows the obtained LCCs as a function of the CI electrode numbers. It can be seen that the Wiener+ACE condition shows the lowest correlation for the lower frequency bands and that Deep ACE shows, in general, the highest LCCs. The results suggest that denoising mid-low frequencies is more challenging, while denoising higher frequencies is easier. This may be due to the predominance of lower-frequency noise signals and the relative scarcity of higher-frequency signals in the target. Specifically, note how LCCs were lower for the SSN noise kind, where low-frequencies are dominant when compared to the ICRA7 noise condition.

**Fig. 4.**
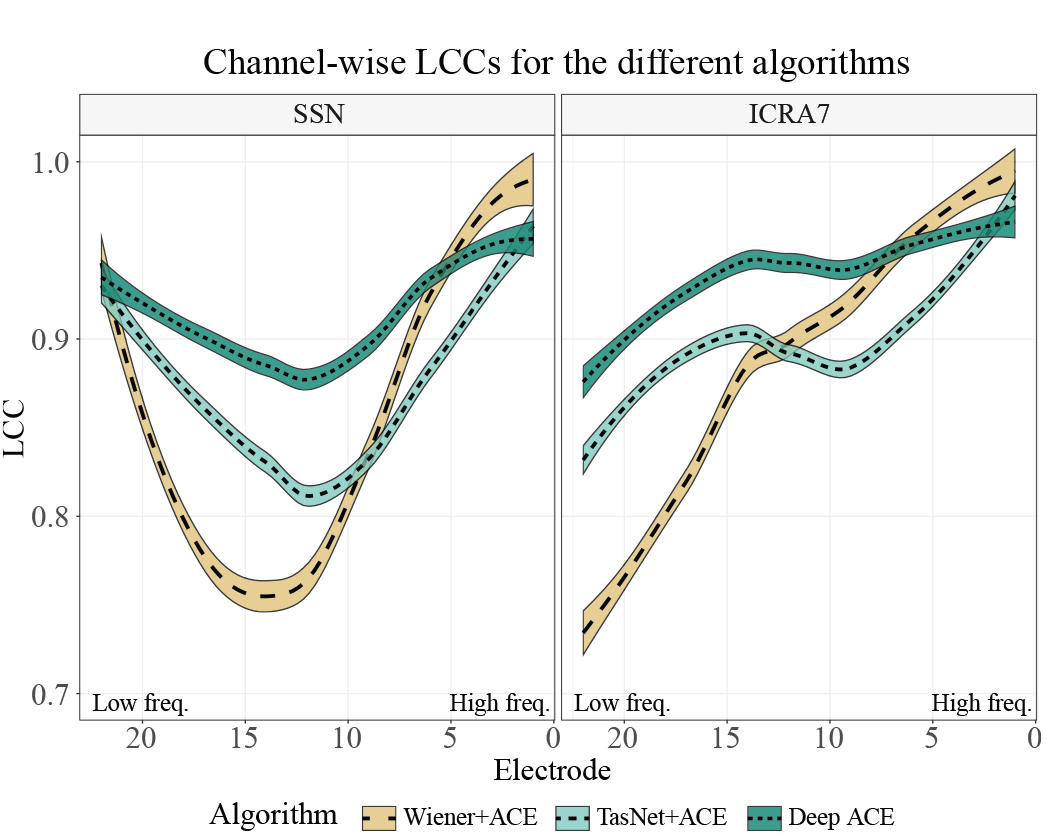
Polynomial regressions showing the channel-wise LCCs between processed and clean electrodograms for the different algorithms and noises using the HSM dataset. Shaded areas represent the 95% confidence level interval [39]. Higher electrode numbers represent lower frequencies.

#### c) VSTOI

Figure 5 illustrates the VSTOI scores obtained by the evaluated algorithms in different speech and noise conditions. In general, the VSTOI scores obtained with the proposed Deep ACE model are higher and agree with the obtained SNRi. These results also represent a substantial improvement compared to the model presented in [20].

**Fig. 5.**
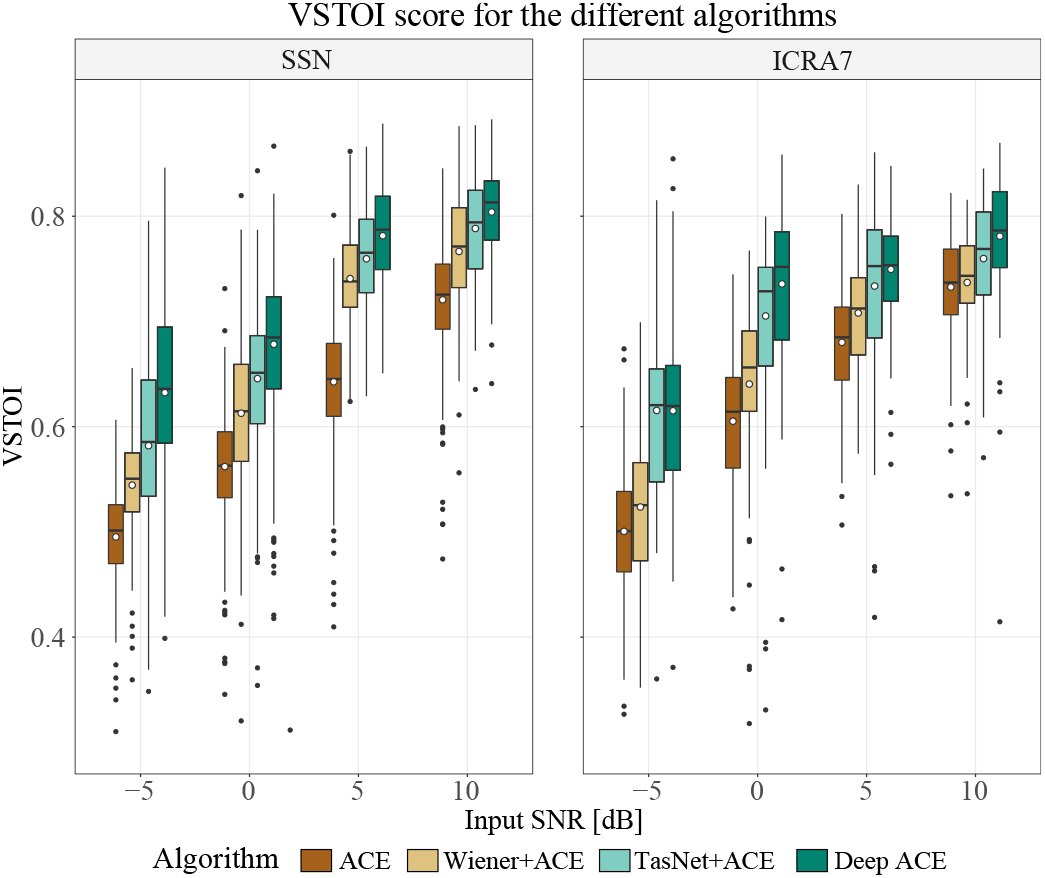
Box plots showing the VSTOI scores for the tested algorithms in SSN and ICRA7 noises for the different SNRs using the HSM speech dataset. All pair-wise differences were statistically significant. The black horizontal bars within each box represent the median for each condition, the circle-shaped marks indicate the mean improvement, and the top and bottom extremes of the boxes indicate the *Q*_3_ = 75% and *Q*_1_ = 25% quartiles, respectively. The box length is given by the interquartile range (IQR), used to define the whiskers that show the variability of the data above the upper and lower quartiles (the upper whisker is given by *Q*_3_ + 1.5·IQR and the lower whisker is given by *Q*_1_ – 1.5·IQR [39]). Black dots indicate observations that fall beyond the whisker range (outliers).

In quiet, the obtained mean VSTOI scores obtained by ACE, TasNet+ACE, and Deep ACE were 0.807, 0.789, and 0.803, respectively.

### 2) Behavioral results

Figure 7 shows the WRS measured in quiet for eight CI subjects. We evaluated ACE and Deep ACE without background noise to test whether the latter introduced any artifacts that compromised the intelligibility of the clean speech signals. A Wilcoxon signed-rank test [40] showed no significant differences between the mean WRS measured using ACE and Deep ACE (*p* = 0.85), confirming that our method coded the clean speech accurately.

**Fig. 6.**
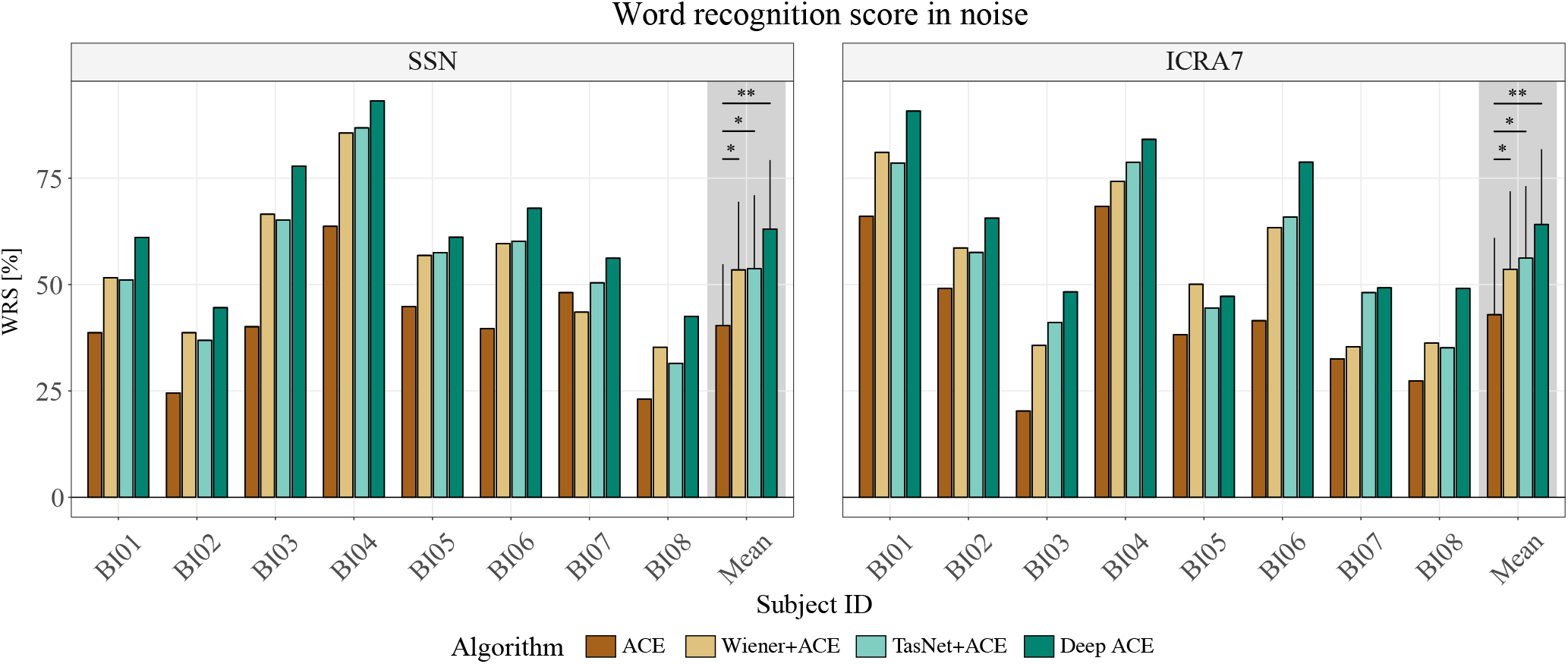
Bar plots showing the mean individual and group mean percentage of correctly understood words by subject for the HSM sentence test in noise for SSN (left panel) and ICRA7 (right panel) noises for all tested conditions. Error bars indicate the standard deviation. Asterisks on top of the significance bar indicate the significance level (* *p*< 0.05, ** *p*< 0.01, *** *p*< 0.001).

**Fig. 7.**
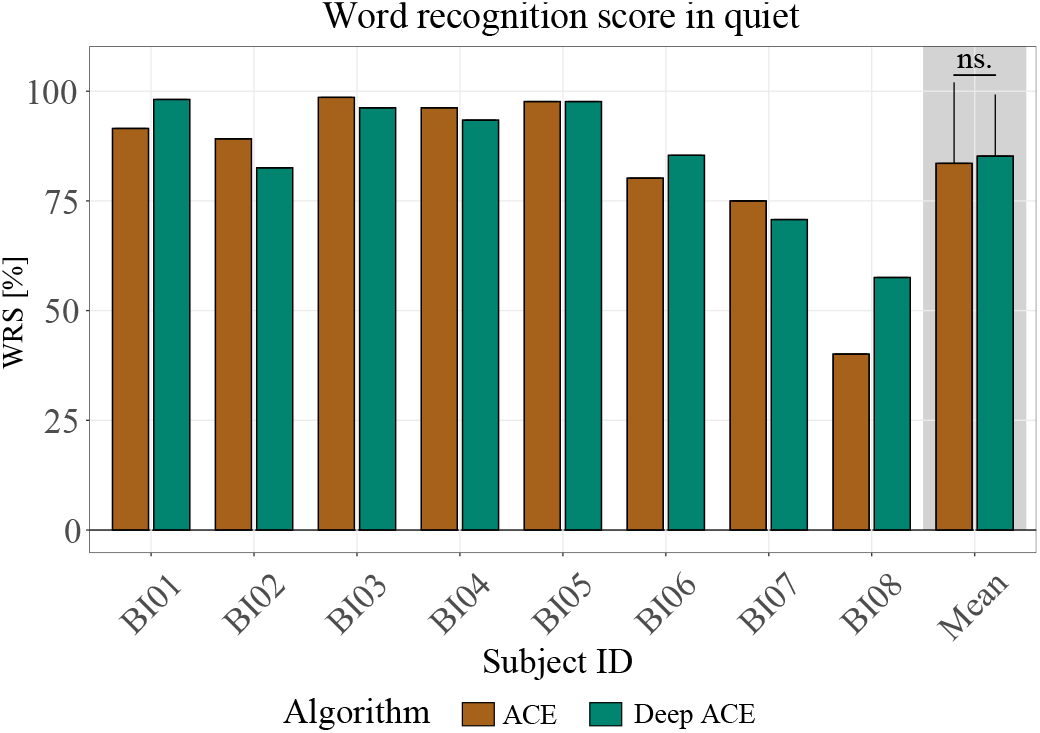
Bar plots showing the mean individual and group mean percentage of correctly understood words by subject for the HSM sentence test in quiet. Error bars indicate the standard deviation.

Figure 6 shows the WRS in noise measured using the different algorithms for the different noises. The normally distributed mean WRS values were evaluated using two 1-way repeated measures analyses of variance (ANOVA; [41]) for each noise condition, with the tested algorithm as the factor. Any ANOVA that revealed a significant effect was followed up by the required post-hoc tests, for which type I error was corrected based on the Holm-Bonferroni method [42].

The ANOVAs revealed a significant effect of algorithm in the measured mean WRS when using SSN background noise [*F*(3, 21) = 3.61, *p* = 0.02] and under ICRA7 background noise [*F*(3, 21) = 2.99, *p* = 0.03]. The ANOVAS were followed up by the corresponding post-hoc tests based on dependent-sample *t*-tests. These revealed that all algorithms outperformed the ACE sound coding strategy in both background noise types. Specifically, for the SSN background noise, ACE obtained a WRS = 40.3%. Wiener+ACE gave a higher benefit in WRS when compared to ACE (*M* = 54.7%, *p* = 0.04), TasNet+ACE also gave higher scores than ACE (*M* = 54.9%, *p* = 0.04), and Deep ACE also outperformed ACE (*M* = 63.1%, *p* = 0.002). For the ICRA7 background noise, ACE obtained WRS = 42.9%. The WRS obtained by ACE was lower than the one measured for Wiener+ACE (*M* = 54.3%, *p* = 0.04), TasNet+ACE (*M* = 56.2%, *p* = 0.03), and Deep ACE (*M* = 64.1%, *p* = 0.007).

In order to assess the WRS benefit obtained with each of the three algorithms, the improvement in WRS with respect to ACE was computed (ΔWRS = WRS_denoised_. – WRS_ACE_). Figure 8 shows the benefit in speech understanding (ΔWRS) provided by the different algorithms for the two noise types. All data were normally distributed so two 1-way repeated measures ANOVA were conducted (one per noise condition) to determine whether the tested algorithms had an impact on the WRS improvement. The ANOVAs yielded a significant effect of algorithm for both SSN background noise [*F*(2, 14) = 3.89, *p* = 0.04] and ICRA7 background noise [*F*(2, 14) = 4.3, *p* = 0.02]. The post-hoc t-tests revealed that the highest improvement was given by Deep ACE for both noise conditions. Deep ACE outperformed Wiener+ACE in SSN (ΔWRS = 22.8%, *p* = 0.02), and in ICRA7 noise (ΔWRS = 21.2%, *p* = 0.005), and it was also superior to TasNet+ACE in SSN (*p* = 0.02), and in ICRA7 noise (*p* = 0.04).

**Fig. 8.**
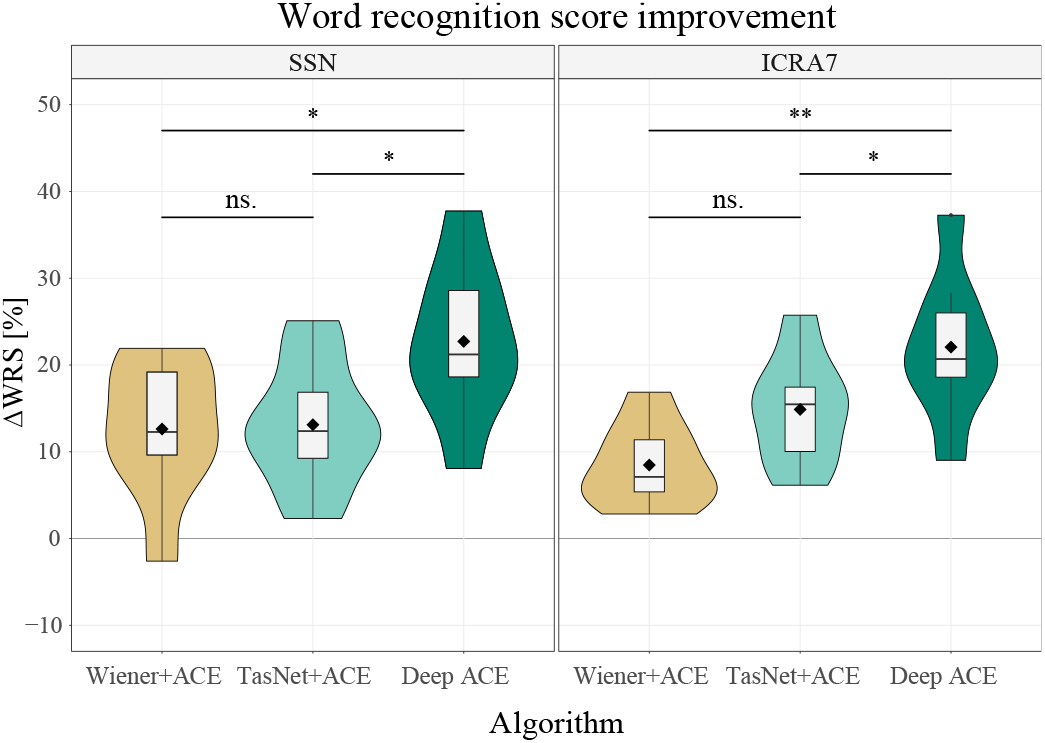
Violin plots showing the WRS improvement by processing the noisy signals with the different algorithms compared to ACE. The black horizontal bars within each of the boxes represent the median for each condition, the diamond-shaped marks indicate the mean improvement, and the top and bottom extremes of the boxes indicate the *Q*_3_=75% and *Q*_1_ = 25% quartiles, respectively. The box length is given by the interquartile range (IQR), used to define the whiskers that show the variability of the data above the upper and lower quartiles (the upper whisker is given by *Q*_3_ + 1.5·IQR, and the lower whisker is given by *Q*_1_ - 1.5·IQR [39]). Asterisks on top of the significance bar indicate the significance level (* *p*< 0.05, ** *p*< 0.01, *** *p*< 0.001). Black dots indicate observations that fall beyond the whisker range (outliers).

## IV. Discussion

In this work, we propose an end-to-end speech coding and denoising strategy for CIs; Deep ACE. The vast majority of speech enhancement algorithms for CIs rely on front-end processing that discards potentially rich sources of information, for this reason, here we investigate an end-to-end deep learning model that merges the denoising preprocessing stage with the CI sound coding strategy. This approach leverages the simplicity of the output signal to be estimated, the electrodogram, which does not require any phase information to be reconstructed, potentially facilitating CI noise reduction.

Combining the noise reduction algorithm with the CI sound coding strategy has the added advantage of reducing processing latency when compared to other front-end methods. For instance, using a front-end TasNet denoising block would result in a latency of 4 ms, whereas the Deep ACE model presented here only introduces 2 ms of latency. This is particularly crucial for devices like CIs that need to transmit signals with minimal latency delays. For example, in the case of single-sided deaf individuals (i.e., CI in one ear and normal hearing in the other), CI processing latency is of utmost importance as these users are exposed to relative sound delay values between the CI and normal hearing ear of 10-12 ms [43]. Here, the goal is to reduce CI processing time to align with the natural delay caused by the traveling wave inside the cochlea, which ranges from 1-9 ms, being longer at lower frequencies [44]. This is desirable because relative delay differences between the CI and acoustic listening sides can disrupt spatial hearing for single-sided CI users [43]. Additionally, lowering latency is important to address any issues with unsynchronization between the speech being spoken and the speech being perceived, and other problems related to audiovisual mismatch that could negatively impact the advantages of lip reading.

This work builds on a previous study [20] which introduced Deep ACE for the first time. Here, we have optimized the architecture and introduced a new CI-specific loss function, aiming at improved speech enhancement performance and greater generalization power. The results indicate that the presented end-to-end CI speech enhancement model outperforms the front-end baseline algorithms in terms of SNRi and predicted speech intelligibility. Additionally, these findings indicate that the model has strong generalization capabilities, performing well with new, unfamiliar data and exhibiting resilience to various types of noise and speech signals, representing a notable advancement over the model featured in [20], which utilized some of the same test materials as those used in the training phase.

The behavioral speech tests with no background noise showed that the proposed end-to-end deep learning coding strategy ‘Deep ACE’ can be used to accurately code the clean speech captured by the CI microphone. Specifically, speech tests in quiet revealed no significant differences in speech understanding between the clinical ACE sound coding strategy and the proposed Deep ACE (see Figure 7). Furthermore, word recognition scores measured in noise showed a benefit of using all the speech-denoising methods, obtaining a statistically significant improvement relative to the baseline ACE condition, as seen in Figure 6. Note that the here observed improvement obtained by the Wiener+ACE using ICRA7 background noise was not observed in [20], however, it is consistent with other studies [9]. This result may be explained by the fact that, in this work, this condition was mostly tested at positive SNRs (see Table III). Finally, when comparing the WRS improvement with respect to ACE obtained by the three tested speech-denoising algorithms, Deep ACE outperformed the other two, obtaining the highest WRS benefit (see Figure 8), this benefit of Deep ACE was also not observed in the listening tests performed in [20]. Although not statistically significant, the TasNet+ACE condition demonstrated a higher WRS improvement score compared to the Wiener+ACE condition when tested with ICRA7 background noise. This outcome is consistent with the objective measures that indicate a greater improvement in SNR and VSTOI scores, as shown in the right panels of Figures 3 and 5.

## V. Conclusions

In this study, we present Deep ACE, a speech coding and denoising sound coding strategy for CIs that utilizes end-to-end deep learning processing. This method aims to provide precise acoustic signal coding like ACE while effectively removing background noise without introducing processing latency. We assessed the performance of the proposed model through both objective measures and listening tests with eight CI users, comparing its performance to the standard ACE and two front-end baseline models, namely the Wiener filter and TasNet. Our results indicated that Deep ACE effectively codes speech signals and outperforms the baseline models in both objective measures and listening tests. These findings suggest that Deep ACE has the potential to replace the current clinical ACE sound coding strategy and improve speech comprehension for CI users in noisy environments.

## Footnotes

1 https://github.com/APGDHZ/DeepACE2.0

## References

[1] J. Wouters, H. J. McDermott et al., “Sound coding in cochlear implants: From electric pulses to hearing,” IEEE Signal Processing Magazine, vol. 32, no. 2, pp. 67–80, 2015.

[2] B. S. Wilson, C. C. Finley et al., “Better speech recognition with cochlear implants,” Nature, vol. 352, no. 6332, pp. 236–238, 1991.

[3] P. Seligman, H. McDermott et al., “Architecture of the spectra 22 speech processor,” Annals of Otology, Rhinology and Laryngology, vol. 104, no. suppl 166, pp. 139–141, 1995.

[4] W. Nogueira, A. Büchner et al., “A psychoacoustic “NofM”-type speech coding strategy for cochlear implants,” EURASIP Journal on Advances in Signal Processing, vol. 2005, no. 18, pp. 1–16, 2005.

[5] I. Hochberg, A. Boothroyd et al., “Effects of noise and noise suppression on speech perception by CI users,” Ear and Hearing, vol. 13, pp. 263–271, 1992.

[6] W. Nogueira, T. Rode et al., “Spectral contrast enhancement improves speech intelligibility in noise for cochlear implants,” The Journal of the Acoustical Society of America, vol. 139, no. 2, pp. 728–739, 2016.

[7] T. Baer, B. C. Moore et al., “Spectral contrast enhancement of speech in noise for listeners with sensorineural hearing impairment: Effects on intelligibility, quality, and response times,” Journal of rehabilitation research and development, vol. 30, pp. 49–49, 1993.

[8] L.-P. Yang and Q.-J. Fu, “Spectral subtraction-based speech enhancement for cochlear implant patients in background noise,” The journal of the Acoustical Society of America, vol. 117, no. 3, pp. 1001–1004, 2005.

[9] N. Guevara, A. Bozorg-Grayeli et al., “The voice track multiband single-channel modified wiener-filter noise reduction system for cochlear implants: patients’ outcomes and subjective appraisal,” International Journal of Audiology, vol. 55, no. 8, pp. 431–438, 2016.

[10] R. Koning, N. Madhu et al., “Ideal time–frequency masking algorithms lead to different speech intelligibility and quality in normal-hearing and cochlear implant listeners,” IEEE Transactions on Biomedical Engineering, vol. 62, no. 1, pp. 331–341, 2014.

[11] Y.-H. Lai, F. Chen et al., “A deep denoising autoencoder approach to improving the intelligibility of vocoded speech in cochlear implant simulation,” IEEE Transactions on Biomedical Engineering, vol. 64, no. 7, pp. 1568–1578, 2016.

[12] Y.-H. Lai, Y. Tsao et al., “Deep learning–based noise reduction approach to improve speech intelligibility for cochlear implant recipients,” Ear and hearing, vol. 39, no. 4, pp. 795–809, 2018.

[13] Y. Hu and P. C. Loizou, “Environment-specific noise suppression for improved speech intelligibility by cochlear implant users,” The Journal of the Acoustical Society of America, vol. 127, no. 6, pp. 3689–3695, 2010.

[14] N. Mamun, S. Khorram et al., “Convolutional neural network-based speech enhancement for cochlear implant recipients,” in Interspeech, vol. 2019. NIH Public Access, 2019, p. 4265.

[15] F. Weninger, H. Erdogan et al., “Speech enhancement with LSTM recurrent neural networks and its application to noise-robust ASR,” in International conference on latent variable analysis and signal separation. Springer, 2015, pp. 91–99.

[16] D. Amodei, S. Ananthanarayanan et al., “Deep speech 2: End-to-end speech recognition in english and mandarin,” in International conference on machine learning, 2016, pp. 173–182.

[17] Y. Luo and N. Mesgarani, “Conv-TasNet: Surpassing ideal time–frequency magnitude masking for speech separation,” IEEE/ACM transactions on audio, speech, and language processing, vol. 27, no. 8, pp. 1256–1266, 2019.

[18] E. C. Cherry, “Some experiments on the recognition of speech, with one and with two ears,” The Journal of the acoustical society of America, vol. 25, no. 5, pp. 975–979, 1953.

[19] F. Bolner, T. Goehring et al., “Speech enhancement based on neural networks applied to cochlear implant coding strategies,” in 2016 IEEE International Conference on Acoustics, Speech and Signal Processing (ICASSP). IEEE, 2016, pp. 6520–6524.

[20] T. Gajecki and W. Nogueira, “An end-to-end deep learning speech coding and denoising strategy for cochlear implants,” in ICASSP 2022-2022 IEEE International Conference on Acoustics, Speech and Signal Processing (ICASSP). IEEE, 2022, pp. 3109–3113.

[21] P. Scalart and J. V. Filho, “Speech enhancement based on a priori signal to noise estimation,” in 1996 IEEE International Conference on Acoustics, Speech, and Signal Processing Conference Proceedings, vol. 2. IEEE, 1996, pp. 629–632.

[22] S. J. Mauger, K. Arora et al., “Cochlear implant optimized noise reduction,” Journal of Neural Engineering, vol. 9, no. 6, p. 065007, 2012.

[23] A. Büchner, M. Brendel et al., “Results of a pilot study with a signal enhancement algorithm for HiRes 120 cochlear implant users,” Otology & neurotology, vol. 31, pp. 1386–90, 2010.

[24] J. R. Hershey, Z. Chen et al., “Deep clustering: Discriminative embeddings for segmentation and separation,” in 2016 IEEE International Conference on Acoustics, Speech and Signal Processing (ICASSP), 2016, pp. 31–35.

[25] D. P. Kingma and J. Ba, “Adam: A method for stochastic optimization,” arXiv preprint arXiv:1412.6980, 2014.

[26] J. L. Roux, S. Wisdom et al., “SDR – half-baked or well done?” in IEEE International Conference on Acoustics, Speech and Signal Processing (ICASSP), 2019, pp. 626–630.

[27] O. U. R. Qazi, B. van Dijk et al., “Understanding the effect of noise on electrical stimulation sequences in cochlear implants and its impact on speech intelligibility,” Hearing Research, vol. 299, pp. 79–87, 2013.

[28] B. Beilharz, X. Sun et al., “LibriVoxDeEn: A corpus for german-to-english speech translation and speech recognition,” in Proceedings of LREC, 2020.

[29] V. Zue, S. Seneff et al., “Speech database development at mit: Timit and beyond,” Speech communication, vol. 9, no. 4, pp. 351–356, 1990.

[30] I. Hochmair-Desoyer, E. Schulz et al., “The hsm sentence test as a tool for evaluating the speech understanding in noise of cochlear implant users.” The American journal of otology, vol. 18, no. 6 Suppl, pp. S83–S83, 1997.

[31] J. Thiemann, N. Ito et al., “DEMAND: a collection of multi-channel recordings of acoustic noise in diverse environmentss,” in Proc. Meetings Acoust, 2013, pp. 1–6.

[32] H. Fastl and E. Zwicker, “Psychoacoustics - facts and models,” Springer, vol. 3rd edition, 2007.

[33] W. A. Dreschler, H. Verschuure et al., “ICRA noises: Artificial noise signals with speech-like spectral and temporal properties for hearing instrument assessment,” Audiology, vol. 40, no. 3, pp. 148–157, 2001.

[34] C. H. Taal, R. C. Hendriks et al., “A short-time objective intelligibility measure for time-frequency weighted noisy speech,” in 2010 IEEE international conference on acoustics, speech and signal processing. IEEE, 2010, pp. 4214–4217.

[35] G. D. Watkins, B. A. Swanson et al., “An evaluation of output signal to noise ratio as a predictor of cochlear implant speech intelligibility,” Ear and Hearing, vol. 39, no. 5, pp. 958–968, 2018.

[36] R. Hinrichs, T. Gajecki et al., “A subjective and objective evaluation of a codec for the electrical stimulation patterns of cochlear implants,” Journal of the Acoustic Society of America, Mar. 2021.

[37] D. Freedman, R. Pisani et al., “Statistics,” Pisani, R. Purves, 4th edn. WW Norton & Company, New York, 2007.

[38] C. H. Taal, R. C. Hendriks et al., “A short-time objective intelligibility measure for time-frequency weighted noisy speech,” in 2010 IEEE international conference on acoustics, speech and signal processing. IEEE, 2010, pp. 4214–4217.

[39] R Core Team, R: A Language and Environment for Statistical Computing, R Foundation for Statistical Computing, Vienna, Austria, 2012, ISBN 3-900051-07-0. [Online]. Available: http://www.R-project.org/

[40] F. Wilcoxon, “Individual comparisons by ranking methods frank,” Biometrics Bulletin, vol. 1, no. 6, pp. 80–83, 1945.

[41] E. R. Girden, ANOVA: Repeated measures. Sage, 1992, no. 84.

[42] O. J. Dunn, “Multiple comparisons among means,” Journal of the American Statistical Association, vol. 56, no. 293, pp. 52–64, 1961.

[43] S. Zirn, J. Angermeier et al., “Reducing the device delay mismatch can improve sound localization in bimodal cochlear implant/hearing-aid users,” Trends in Hearing, vol. 23, 2019.

[44] M. A. Ruggero and A. N. Temchin, “Similarity of traveling-wave delays in the hearing organs of humans and other tetrapods,” Journal for the Association for Research in Otolaryngology, vol. 8, no. 2, pp. 153–166, 2007.

